# SPLICE-q: a Python tool for genome-wide quantification of splicing efficiency

**DOI:** 10.1101/2020.10.12.318808

**Authors:** Verônica R Melo Costa, Julianus Pfeuffer, Annita Louloupi, Ulf A V Ørom, Rosario M Piro

## Abstract

**Background:** Introns are generally removed from primary transcripts to form mature RNA molecules in a post-transcriptional process called splicing. An efficient splicing of primary transcripts is an essential step in gene expression and its misregulation is related to numerous human diseases. Thus, to better understand the dynamics of this process and the perturbations that might be caused by aberrant transcript processing it is important to quantify splicing efficiency.

**Results:** Here, we introduce SPLICE-q, a fast and user-friendly Python tool for genome-wide **SPLIC**ing **E**fficiency **q**uantification. It supports studies focusing on the implications of splicing efficiency in transcript processing dynamics. SPLICE-q uses aligned reads from strand-specific RNA-seq to quantify splicing efficiency for each intron individually and allows the user to select different levels of restrictiveness concerning the introns’ overlap with other genomic elements such as exons of other genes. We applied SPLICE-q to globally assess the dynamics of intron excision in yeast and human nascent RNA-seq. We also show its application using total RNA-seq from a patient-matched prostate cancer sample.

**Conclusions:** Our analyses illustrate that SPLICE-q is suitable to detect a progressive increase of splicing efficiency throughout a time course of nascent RNA-seq and it might be useful when it comes to understanding cancer progression beyond mere gene expression levels. SPLICE-q is available at: https://github.com/vrmelo/SPLICE-q

## 1. Background

Eukaryotic genes are mostly composed of a series of exons intercalated by sequences with no coding potential called introns. These sequences are generally removed from primary transcripts by a post-transcriptional process called splicing to form mature RNA molecules. This highly regulated process consists basically of a series of hydrolysis and ligation reactions led by the spliceosome [1]. The exon-intron boundaries, i.e., the splice junctions, together with the branch point, a short sequence located 18-40 nucleotides upstream of the intron’s 3’ splice junction [2] and the polypyrimidine tract [3], are recognized by the spliceosome. These events promote the correct folding necessary for the intron’s excision and are followed by the correct pairing of the exon-exon boundaries. In metazoans, further sequences are required for recruiting different trans-acting regulatory factors. These will act as spliceosome regulators as well as splice site choice modulators and are particularly important for efficient transcript processing [4].

Splicing is dynamic and occurs mostly during or immediately after the transcription of a complete intron. Co-transcriptional splicing was first suggested in *D. melanogaster* chorion genes using electron microscopy to observe the assembly of spliceosomes at the splice junctions in nascent transcripts [5]. More recently, genome-wide studies in different cell lines and organisms using nascent RNA showed introns being spliced shortly after their transcription is finished: in *S. cerevisiae*, data revealed polymerase pausing within the terminal exon, permitting enough time for splicing to happen before release of the mature RNA [6]; and nascent RNA also indicated that most introns in *D. melanogaster* are co-transcriptionally spliced [7], as well as in mouse [8] and many human cells and tissues [9–11].

Splicing is an essential step in gene expression and its misregulation is related to numerous human diseases [12–15]. Up to 15% of mutations that cause genetic disease have been suggested to affect pre-mRNA splicing [16]. Thus, to better understand the dynamics of splicing and the perturbations that might be caused by aberrant transcript processing, it is important to quantify splicing efficiency. The efficiency of splicing is commonly quantified by means of RT-qPCR with primers that span exon-exon and exon-intron boundaries [17]. Yet, this methodology can only investigate a limited number of genes. By contrast, transcriptomics technologies, such as RNA-Seq, allow these analyses from a genome-wide point of view. One interesting approach to globally determine splicing efficiencies is to assess nascent transcripts within short intervals after the transcription has started. Experimentally, this can be achieved through metabolic labeling or purification of chromatin-associated nascent RNAs.

For intron-containing transcripts, splicing efficiency can be determined with different frameworks that use read counts on intronic and exonic regions. Short-read RNA-Seq is currently the main approach using either nascent or total RNA. Conceptually, splicing efficiency can be observed either from an intron-centric point of view—to investigate whether an intron has been spliced out—or from an exon-centric point of view—to investigate whether an exon has been correctly spliced within the context of its transcript.

Khodor et al. [7] used an intron-centric method to estimate the unspliced fraction of introns in *D. melanogaster* by taking the ratio of the read coverage of the last 25 bp of an intron and the first 25 bp of the following exon. In this way, introns where the RNA polymerase has not yet reached the acceptor splice site are not included but the metric is not guaranteed to take values between 0 and 1 and does hence not constitute an efficiency metric in the strict sense. Tilgner et al. [10] used deep-sequencing of human subcellular fractions and developed an exon-centric “completed splicing index” (coSI) which takes reads spanning the 5’ and the 3’ splice junctions of an exon and computes the fraction of reads indicating completed splicing, i.e., which span from exon to exon, to study co-transcriptional splicing.

By explicitly considering also reads which span from the upstream exon directly to the downstream exon, this approach includes exon skipping events, but coSI values for the first and last exon of a transcript cannot be determined. More recently, Převorovský et al. [18] presented a workflow for genome-wide determination of intron-centric splicing efficiency in yeast. The efficiencies are quantified for the 5’ and 3’ splice junctions separately as the number of “*transreads*” (split reads spanning from exon to exon) divided by the number of reads covering the first or last base of the intron, respectively. Although the authors call their metric “splicing efficiency”, it is not limited to a range from 0 to 1 and it is not clear how cases without intronic reads (divisions by zero) are handled. Other drawbacks of this workflow are that it consists of numerous open-source tools and custom shell and R scripts and that it was explicitly developed for yeast.

Although the above-mentioned frameworks for calculating splicing efficiency from RNA-seq data exist, there is more to add to their respective limitations. The bioinformatics steps involved might be challenging—including difficulties in running workflows that require long running times and the installation of numerous tools—especially for experimental biologists. Thus, here we introduce SPLICE-q, a user-friendly open-source Python tool for genome-wide **SPLIC**ing **E**fficiency **q**uantification from RNA-seq data. SPLICE-q quantifies splicing efficiency for each intron individually and allows the user to select different levels of restrictiveness concerning an intron’s overlap with other genomic elements. We show the usefulness of SPLICE-q by applying it to time-series nascent and steady-state RNA-seq data from human and yeast.

## 2. Implementation

### 2.1. SPLICE-q workflow and parameters

SPLICE-q is a tool, implemented in Python 3, for quantification of individual intron splicing efficiencies from strand-specific RNA-seq data. SPLICE-q’s main quantification method uses splicing reads—both split and unsplit—spanning the splice junctions of a given intron (**Fig. 1**). Split reads are junction reads spanning from one exon to another, thus indicating processed transcripts from which the individual intron has already been excised. Intuitively, unsplit reads are those spanning the intron-exon boundaries (covering both sides of the splice junction), hence, indicating transcripts from which the intron has not yet been spliced out. As an alternative measure for splicing efficiency, SPLICE-q computes an inverse intron expression ratio, which compares the introns’ expression levels with those of their flanking exons.

**Fig. 1:**
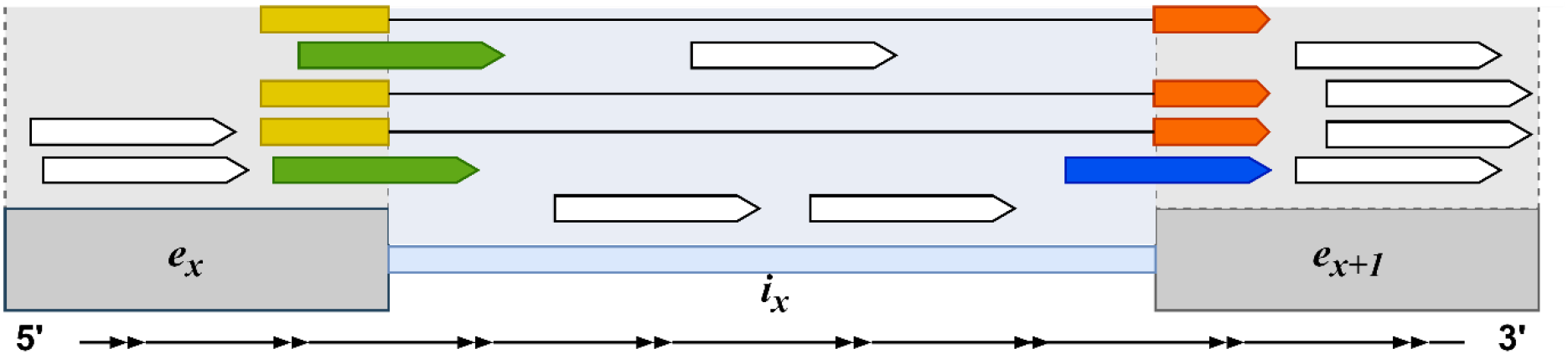
Read assignment scheme for splicing efficiency (SE) and inverse intron expression ratio (IER). Illustration of the reads used by SPLICE-q to quantify SE and IER. In yellow, split reads at the 5’ splice junction; in orange, split reads at the 3’ splice junction; in green, unsplit reads at the 5’ splice junction; in dark blue, unsplit reads at the 3’ splice junction. In gray and blue, the areas covering the exons and introns, respectively. In white, reads not overlapping splice junctions.

SPLICE-q is also sensitive to the overlap of genomic elements. In other words, SPLICE-q takes into consideration when a genome shows overlapping features that can cause issues with a correct assignment of reads to specific introns or exons. For example, for intron-exon boundaries overlapping exons of other genes, seemingly unsplit reads might instead stem from exonic regions of the overlapping genes. This is problematic due to the RNA-seq methodology’s limitation that makes it difficult to confidently determine without ambiguity to which genomic element, exon or intron, these reads should be attributed [19].

Therefore, SPLICE-q allows the user to select different levels of restrictiveness for strand-specific filtering, including (i) Level 1: keep all introns in the genome regardless of overlaps with other genomic elements; (ii) Level 2: select only introns whose splice junctions do not overlap any exon of a different gene; (iii) Level 3: select only introns that do not overlap with any exon of the same or a different gene (**Fig. 2**). Other filters, including the minimum read coverage at splice junctions, can also be set up according to users’ necessities (Additional file 1: Table 1).

The two necessary input files are:

i. A Binary Alignment Map (BAM) file with RNA-seq reads aligned to the reference genome.
ii. A genome annotation file provided by GENCODE [20] or Ensembl [21] in Gene Transfer Format (GTF) containing information on exons and the genes and transcripts they are associated with.

**Fig. 2:**
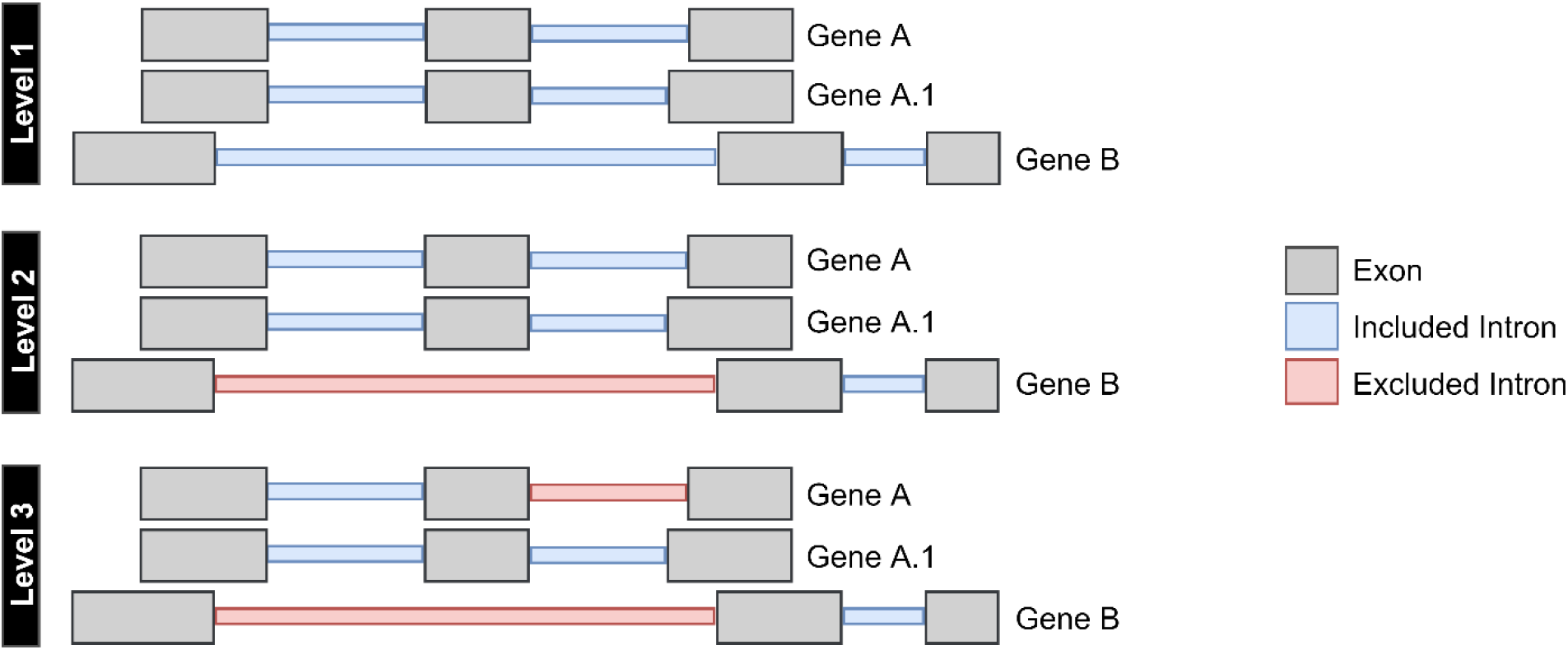
SPLICE-q’s levels of restrictiveness. (Level 1) keep all introns in the genome regardless of overlaps with other genomic elements; (Level 2) select only introns whose splice junctions do not overlap any exon of a different gene; (Level 3) select only introns that do not overlap with any exon of the same or a different gene. A and A.1 are isoforms of the same gene (A) and B represents a different gene.

SPLICE-q’s internal default workflow comprises of the following major steps (**Fig. 3**):

i. Parsing of genomic features from the GTF file;
ii. Locating and annotating introns and splice junctions from the GTF file’s exon coordinates;
iii. Filtering of introns according to the level of restrictiveness based on the overlap of genomic elements;
iv. Selection of split and unsplit reads at the splice junctions according to the reads’ concise idiosyncratic gapped alignment report (CIGAR), and subsequent coverage calculation for each splice junction.
v. Computation of splicing efficiencies. SPLICE-q parses the exon-centric GTF file and infers the corresponding intron coordinates, partially adapting related functions of GTFtools [22]. For Level 3 filtering, when the user chooses to include the inverse intron expression ratio, the workflow includes two additional steps (Additional file 1: Fig. S1):
vi. Computation of median per-base coverages of introns and their flanking exons
vii. Computation of the inverse intron expression ratios.

**Fig. 3:**
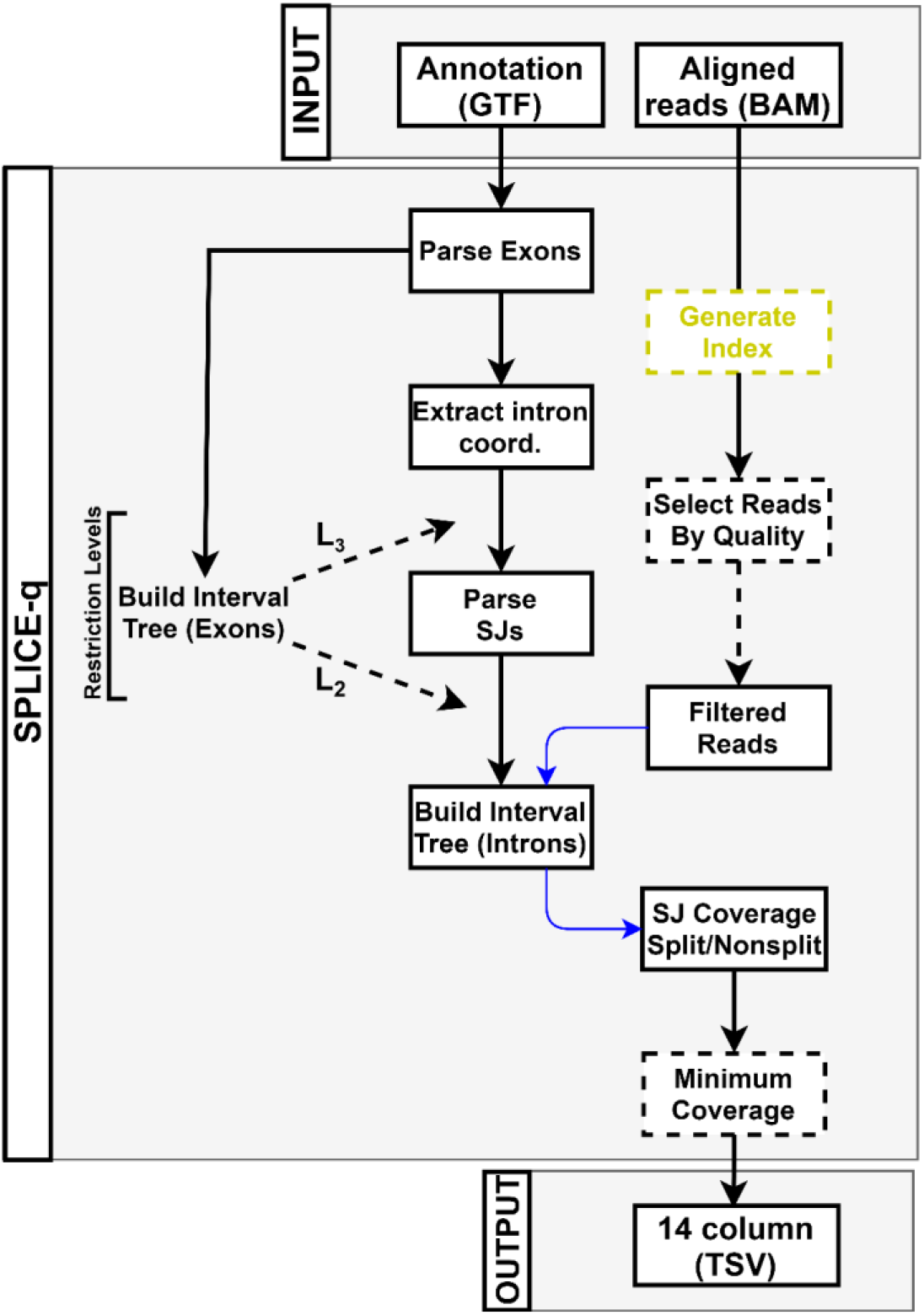
SPLICE-q’s default workflow. Dashed lines indicated steps which depend on parameter settings. Solid lines represent the mandatory steps of the workflow. A BAM index (.bai) file is generated if not provided by the user (yellow). Arrows in blue represent a lookup in the data structure they pass through. SJ = splice junction; TSV = tab-separated values.

### 2.2 Quantifying splicing efficiency and inverse intron expression ratio

#### Splicing efficiency (*SE*)

SPLICE-q uses split and unsplit junction reads to quantify *SE* for each intron individually. It determines the RNA-seq reads mapping to both splice junctions of each given intron *i*, distinguishes split (*S*) and unsplit (*N*) reads for the 5’ and 3’ splice junctions and estimates a splicing efficiency score (*SE_i_*) as a function of the corresponding read counts as follows:

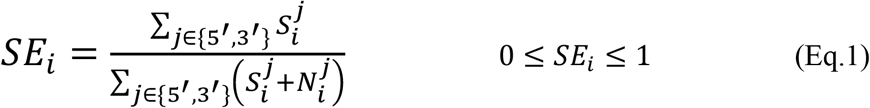

An *SE* of 0 indicates that the intron has not been spliced out in any of the transcripts from which the junction reads originate, which may be due to late splicing in case of nascent RNA-seq or intron retention in case of steady-state RNA-seq. An *SE* of 1 means completed splicing on all transcripts. Therefore, *SE* values ranging between 0 and 1 approximate the fraction of molecules which have already been spliced. This quantification method makes it possible to compare spliced and unspliced intron rates directly.

#### Inverse intron expression ratio (*IER*)

as an alternative measure for splicing efficiency when using Level 3 filtering, SPLICE-q also provides the inverse of the ratio of intron expression to exon expression, where *I_x_* is the median per-base read coverage of the *x*-th intron of a given transcript and *E_x_* and *E*_*x*+1_ represent the corresponding median coverages of the flanking exons:

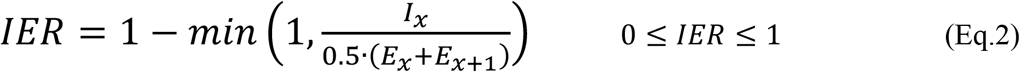

Here, the focus lies specifically on the per-base median coverage of all reads mapping to the involved genomic elements (exonic and intronic reads) rather than just the splice junctions (**Fig. 1**). As explained above, a high *SE* indicates that an intron was spliced out of a large fraction of transcripts. This scenario should display high read coverage in the exons and low coverage or none in the intron. In other words, peaks of mapped reads are observed in the surrounding exons when compared to the intron itself. On the contrary, introns with a low *SE* should have read coverage profiles more similar to the surrounding exons.

## 3. Results and Discussion

### 3.1. Fast and user-friendly quantification of splicing efficiency

Due to its simplicity and efficient data structure for working with genomic intervals, SPLICE-q’s run time with default parameters for approximately 100 million input reads mapped to the human genome is 18 minutes using a MacBook Pro with a dual-core Intel Core i5 processor and 8GB of RAM. By increasing the number of processes to 4 or 8, which is not an issue considering nowadays’ number of processor cores of most laptops and desktops, the running time on an AMD Opteron 6282 SE with 516GB of memory is less than 2 minutes (Additional file 1: **Fig. S2a**). Memory usage is low, being approximately that of the GTF file size (1.4 GB for the human genome; Additional file 1: **Fig. S2b**). SPLICE-q’s approach provides major advantages over previous workflows which may require the installation of numerous tools and suffer from long running times.

### 3.2. Splicing kinetics in human and yeast

We applied SPLICE-q to globally assess the kinetics of intron excision. The goal here is to show the tool’s applicability using different data. For this purpose, we performed three different analyses using data from two species and different methodologies (Additional file 1: Materials and Methods). The first time-series sequencing dataset consists of BrU-labeled HEK293 cells with 15 minutes pulse labeling of nascent RNA and subsequent sequencing of labeled RNA after 0, 15, 30, and 60 minutes (pulse-chase) [23]. On average we obtained ~200 million reads per sample, ~85% of which were uniquely mapped. The nascent RNA samples were compared to an unlabeled steady-state control of the same cell line [24]. SPLICE-q was applied with default parameters: filtering level 3, a minimum coverage of 10 uniquely mapped reads at each splice junction, and a minimum intron length of 30 nucleotides [25]. Only introns passing the filters in all samples after running SPLICE-q were kept, totalizing 13,178 introns. As expected, SPLICE-q detects a progressive increase of *SE* throughout the time course (**Fig. 4a**). Interestingly, at 0 and 15 minutes, *SE* scores are already high with a median of 0.71 and 0.75, respectively. This agrees with previous studies showing that splicing is predominantly co-transcriptional in humans and for the most part happens immediately after the transcription of an intron is completed, when the RNA polymerase has proceeded only a few bases into the downstream exon [5, 6, 9–11]. However, the results also illustrate that even 60 minutes after the pulse-labeling of newly synthesized RNA, there is a significantly larger fraction of introns which have not yet been excised from the transcripts than in the steady-state control.

**Fig. 4:**
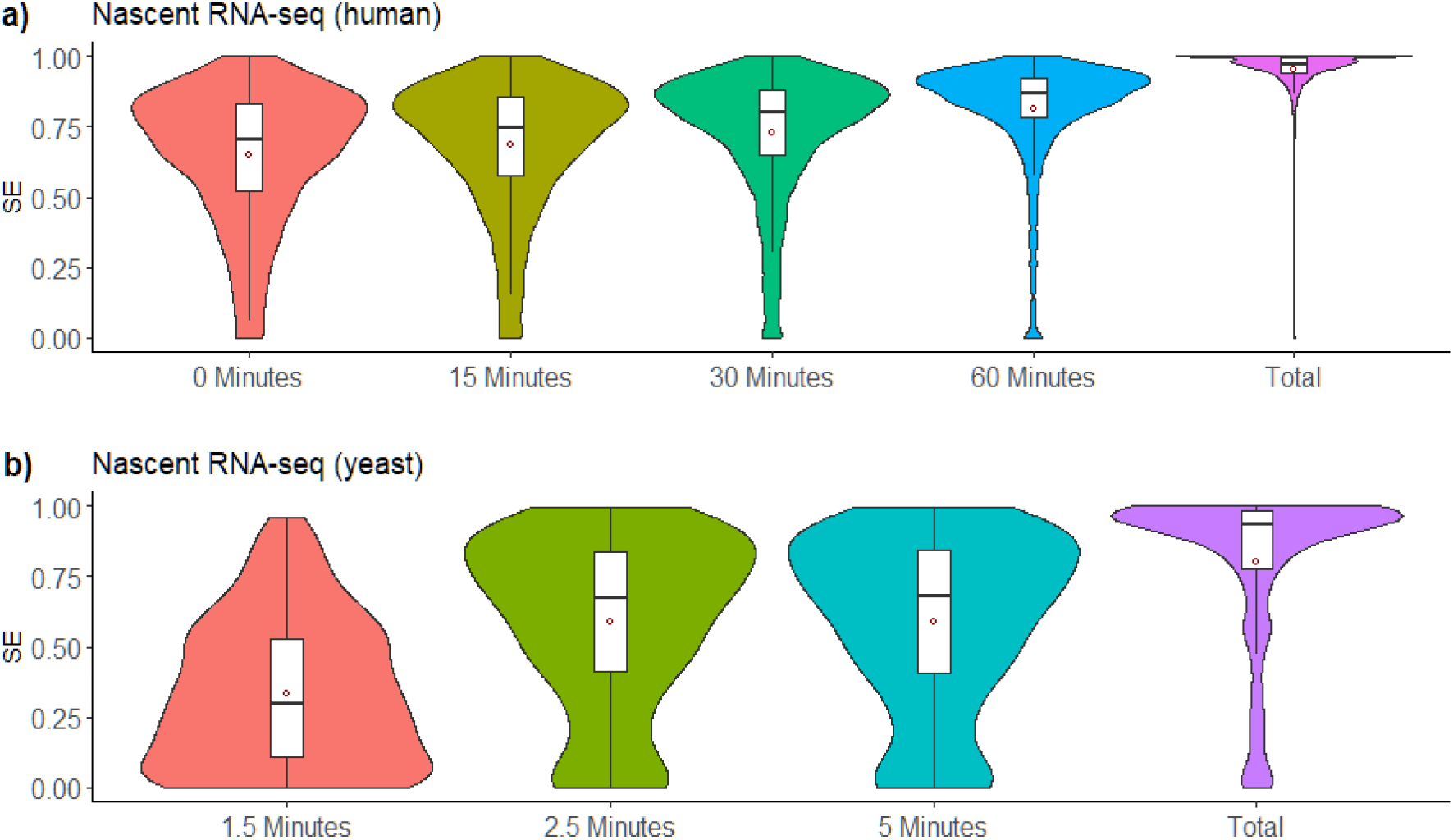
Splicing kinetics using different datasets. **a)** Time-series nascent and steady-state (total) RNA-seq of labeled HEK293 cells; **b)** Time-series nascent and steady-state RNA-seq of *S. cerevisiae*.

We chose a second dataset [26] which would allow us to quantify splicing efficiency of nascent RNA within a finer time scale. These sequencing experiments were performed with 4-thiouracil labeled RNA (4tU-seq) from *Saccharomyces cerevisiae*. Nascent RNA was labeled for an extremely short time (1.5, 2.5 and 5 minutes) and then sequenced (**Fig. 4b**). Unlabeled control samples were also generated. After alignment of the raw data, we obtained an average of over 50 million uniquely mapped reads per sample and 246 introns shared between all samples after running SPLICE-q with the above-mentioned default parameters and filtering level 2. The *SE* at 1.5 minutes has a median of 0.29 while, strikingly, there is an increase of 131% in just one minute, with a median *SE* of 0.67 at 2.5 minutes. This value does not alter in the next time point and the unlabeled control sample shows a median *SE* of 0.93. This brief analysis suggests how essential it is to perform short labeling in *S. cerevisiae* in order to assess its splicing kinetics since some transcripts approximate steady-state levels in a time as short as 2.5 minutes (**Fig. 4b**).

Lastly, we show how SPLICE-q can also be applied to quantify intron retention in steady-state RNA-seq data. For this purpose, we used data coming from a prostate cancer sample along with its matched normal tissue (patient 15 of ref. [27]). Since for each of the tissues two replicates were available, we computed splicing efficiencies for each replicate and then averaged the results for the tumor tissue and the normal tissue.

Prostate cancer is one of the most common cancer types in men [28]. SPLICE-q detected relatively high splicing efficiencies—median *SE* of 0.96 in both the tumor and the normal sample—in the 66,389 introns shared across the sample pair after running the tool with default parameters. This is expected when the tool is applied to steady-state RNA-seq data. Although this overview suggests that there is no alteration in average splicing efficiency levels between normal and tumor tissue, a closer look showed interesting changes for individual introns. One intriguing example is *Prostate cancer associated 3* (PCA3), a long noncoding RNA highly expressed in prostate cancer and widely known as a prostate cancer-specific biomarker of high specificity [29]. It has been found to be involved in the proliferation and survival of prostate cancer cells by multiple mechanisms, including the modulation of androgen receptor signaling, the inhibition of the tumor suppressor PRUNE2, and possibly by acting as a competing endogenous RNA (ceRNA) for *High mobility group box 1* (HMGB1) via sponging of *miR-218-5p* [29–31]. Interestingly, PCA3’s second intron located at chr9:76,782,833-76,783,704 has an *SE* of 0.57 in normal tissue and a much higher *SE* of 0.90 in the tumor (**Fig. 5a**), suggesting that PCA3 might not only be overexpressed but also more efficiently spliced.

**Fig. 5:**
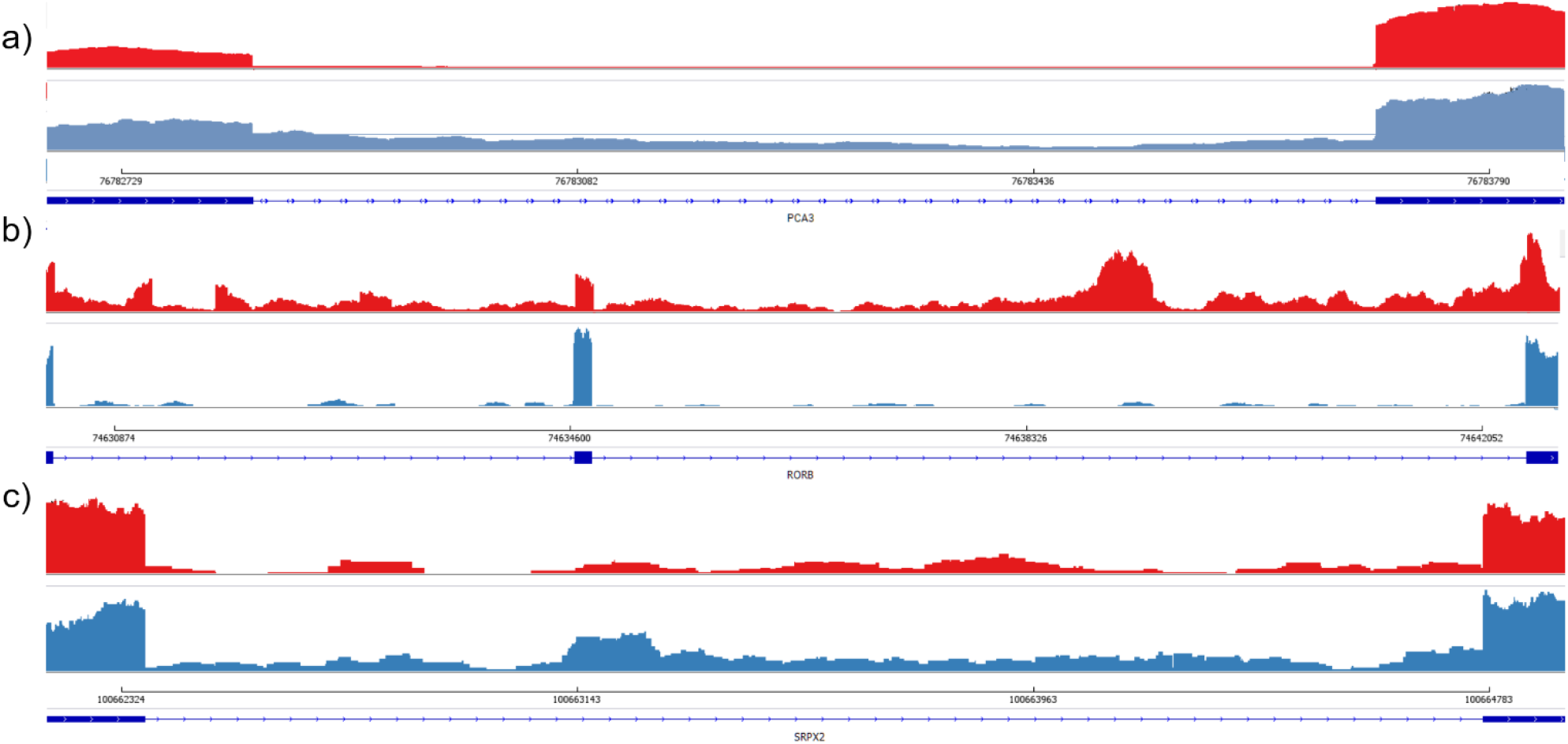
Read coverage of selected introns in the prostate cancer and the normal control sample. IGV views of representative cases of introns from different genes comparing prostate cancer vs. normal samples. a) Intron located at chr9:76,782,833-76,783,704 of PCA3; **b)** Introns located at chr9:74,630,368-74,634,630 and chr9:74,634,773-74,642,413 of RORB; and **c)** Intron located at chrX:100,662,368-100,664,773 of SRPX2. Tumor and normal samples are represented in red and blue, respectively.

Variation in splicing efficiency can be also observed among protein coding genes. The *retinoic acid-related orphan receptor β* (RORβ, encoded by the gene RORB) was recently reported to inhibit tumorigenesis in colorectal cancer *in vivo*. When RORβ was overexpressed, tumorigenic capacity of the cells was significantly reduced, suggesting that this protein acts as a tumor suppressor in colorectal cancer [32]. Intriguingly, we found two of the RORB introns—located at chr9:74,630,368-74,634,630 and chr9:74,634,773-74,642,413—to have reduced splicing efficiencies in the prostate cancer sample (*SEs* of 0.99 and 0.98 in the normal control and 0.63 and 0.60 in the tumor, respectively) (**Fig. 5b**). Contrasting, *Sushi repeat-containing protein X-linked 2*, or simply SRPX2, shows the opposite splicing efficiency profile with an intron at the coordinates chrX:100,662,368-100,664,773 being less efficiently spliced in the control sample (*SE* of 0.59) than in the tumor (*SE* of 0.90, **Fig. 5c**). Previous studies showed SRPX2 to play an important role in cancer development and progression. In colorectal cancer, the overexpression of SRPX2 may promote invasiveness of tumor cells [33], and in prostate cancer, a knockdown of SRPX2 affected the proliferation, migration and invasion of cancer cells by partially suppressing the PI3K/Akt/mTOR signaling pathway [34]. PI3K/Akt/mTOR regulates cell proliferation and survival in different cancer types and is usually activated in advanced prostate cancer [35, 36]. Furthermore, the suppression of this signaling pathway was reported to reduce cell motility and invasion in prostate cancer [37].

These examples illustrate that gene regulation may go beyond the mere expression levels, with a gain or loss of splicing efficiency potentially acting as a superposed mechanism that may be beneficial to tumor development.

## 4. Conclusions

Here we introduced SPLICE-q, an efficient and user-friendly tool for splicing efficiency quantification. SPLICE-q enables the quantification of splicing through two different methods (*SE* and *IER*) and is sensitive to the overlap of genomic elements. We demonstrated SPLICE-q’s usefulness showing three use cases, including two different species and experimental methodologies. Our analyses illustrate that SPLICE-q is suitable to detect a progressive increase of splicing efficiency throughout a time course of strand-specific nascent RNA-seq data. Likewise, SPLICE-q can be applied to strand-specific steady-state RNA-seq data and might be useful when it comes to understanding cancer progression beyond mere gene expression levels.

## Availability and Requirements

**Project name:** SPLICE-q

**Project homepage:** https://github.com/vrmelo/SPLICE-q

**Operating system (s):** Linux, macOS, and Windows 10 Subsystem for Linux.

**Programming language:** Python 3

**Other requirements:** Python 3.x., including packages PySam and InterLap.

**License:** GPL-2

**Any restrictions to use by non-academics:** None

## Supplementary information

**Additional file 1:** Supplementary file contains additional figures, tables and Materials and Methods.

## Acknowledgments

The authors thank A. Marsico, A. Mayer and M. Vingron for fruitful discussions and ideas on this work and related topics. Many thanks to E. Ntini for her support, patience and ideas, especially in the initial steps of this work.

## Authors contributions

VRMC, UAO and RMP designed the study. AL performed the pulse-chase experiments on HEK293 cells. VRMC and JP implemented the method. VRMC analyzed the data. VRMC and RMP wrote the paper.

## Declarations

### • Ethics approval and consent to participate

Not applicable.

### • Consent for publication

Not applicable.

### • Availability of data and materials

The datasets generated and/or analyzed during the current study can be downloaded from the NCBI Gene Expression Omnibus repository with accession numbers GSE92565, GSE83561, GSE84722, GSE70378 and GSE133626.

