## Additional file 1 for "SPLICE-q: a Python tool for genome-wide quantification of splicing efficiency"

#### Supplementary Tables and Figures

**Table 1: Summary table of parameters**.

| Parameter | Description |
|---|---|
| ***MinCoverage*** | Minimum number of reads spanning each splice junction (Default = 10). |
| ***MinReadQuality*** | Mapping quality. By default, only uniquely mapped reads are included (Default = 10). |
| ***MinIntronLength*** | Minimum intron length. Default value is optimal for analysis using human RNA-seq data (Default = 30) |
| ***ChromsList*** | List of chromosome names (Default: chr1-720, I-XVI, 2L, 2R, 3L, 3R, Z, W.) |
| ***FilterLevel*** | (1) keep all introns in the genome regardless of overlaps with other genomic elements.<br>(2) select only introns whose splice junctions do not overlap any exon in different genes<br>(3) select only introns that do not overlap with any exon of the same or different gene (Default). |
| ***IERatio*** | Running mode that additionally outputs the Inverse Intron Expression Ratio (IER). Requires *FilterLevel* 3. |
| ***NProcesses*** | Multiple concurrent processes are used to minimize running times and the number of processes can be adjusted by the user through this parameter. |

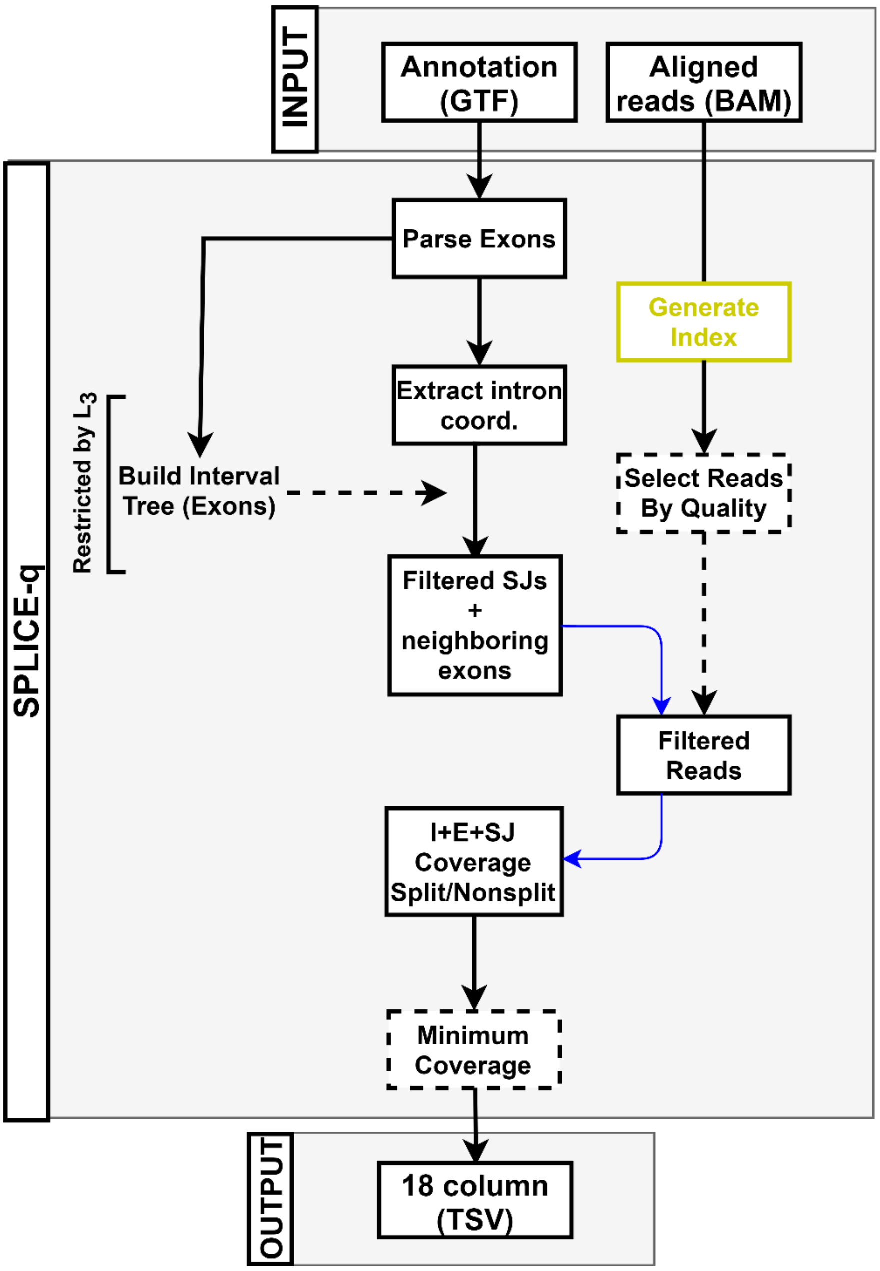


**Fig. S1: SPLICE-q's inverse intron expression ratio (IER) workflow.** Dashed lines indicated steps which depend on parameter settings. Solid lines represent the mandatory steps of the workflow. Arrows in blue represent a lookup in the data structure they pass through. I = intron; E= exon; SJ = splice junction; TSV = tab-separated values.

**a)**

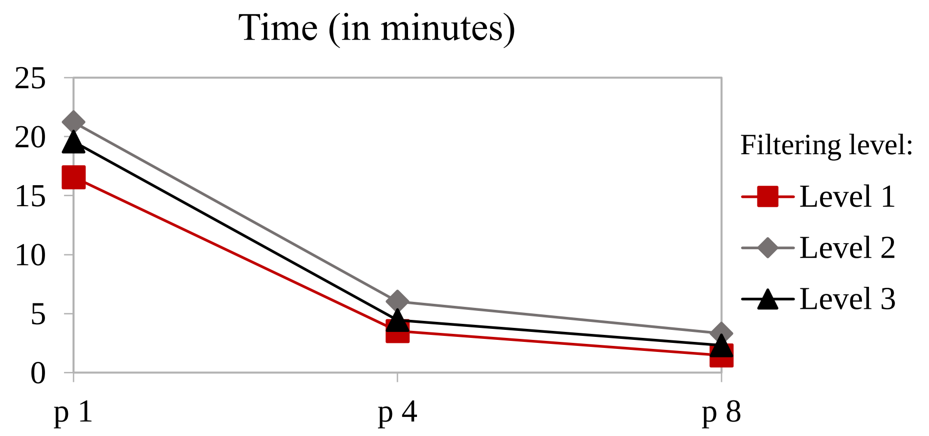


**b)**

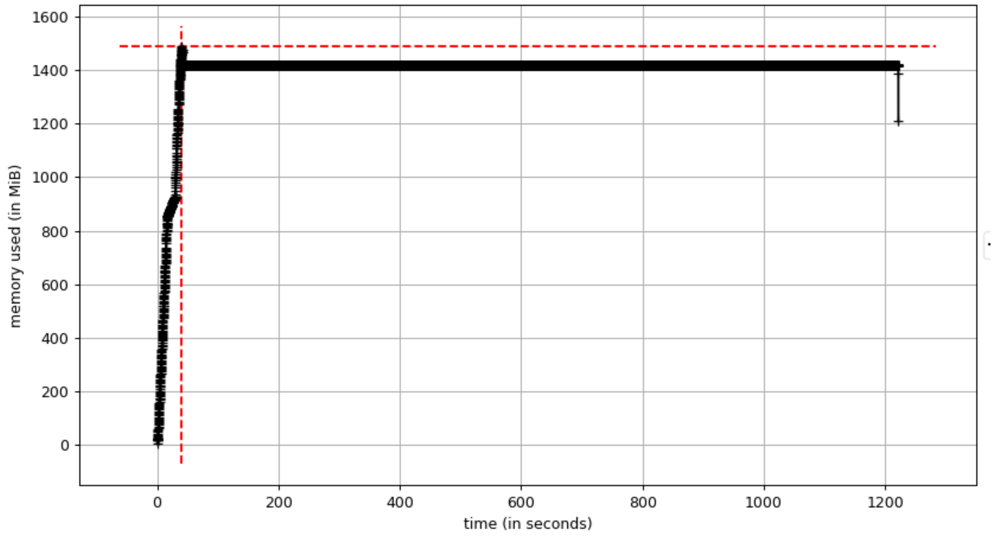


**Figure S2**: **SPLICE-q's run time and memory usage**. **a)** Run time for approximately 100 million input reads mapped to the human genome (Linux, 64x AMD Opteron 6282 SE, 516GB). **b)** Memory usage for 1.4GB GTF. Time in seconds. p = Number of processes (*NProcesses*).

### Materials and Methods

#### Pulse-chase essay, RNA-seq and read mapping

Human embryonic kidney cells (HEK293) cells were incubated for 15 minutes with 2mM of 5-bromouridine (BrU, pulse). Then, the cells were either collected immediately (0 minutes) or chased for 15, 30 and 60 minutes prior to RNA purification and selection of BrU-labeled RNA as described in [1]. The sequencing library was prepared with the TrueSeq Stranded Total RNA Kit (Illumina). Sequencing was performed in triplicate on the Illumina HiSeq 2500 platform to obtain an average of ~200 million reads per sample. Replicates read coverage are highly correlated with an average $\rho = 0.95$ which satisfies the ENCODE consortium recommendations for biological replicates [2]. The strand-specific reads were mapped to the human reference genome *GRCh38.p10* with STAR v2.7.1a [3] according to recommendations from the STAR manual 2.4.0.1.  The genome index for STAR was built on the genome annotation from GENCODE v27[1]. An average of ~85% of the reads in all samples were uniquely mapped. The GEO [4] accession numbers for these sequencing data are GSE92565, GSE83561 and GSE84722.

1 ftp://ftp.ebi.ac.uk/pub/databases/gencode/Gencode_human/release27/gencode.v27.annot ation.gtf.gz

#### Other datasets

The other datasets processed and analyzed are described below.

**Table 1: Datasets used in the study.**

| Accession/ Reference | Genome/ Annotation | Description |
|---|---|---|
| GSE84722 [6] | GRCh38.p10/ gencode v27 | Total RNA-seq of HEK293 cells. Sequenced on HiSeq2500. |
| GSE70378 [7] | Ensembl R64-1-1 | *S. cerevisiae* labeled with 4tU labeling for 1.5, 2.5 and 5 minutes. Total RNA-seq also performed. All experiments were performed in triplicate. Sequenced on HiSeq2500. |
| GSE133626 [8] | GRCh38.p10/ gencode v27 | Total RNA from fresh frozen prostate cancer tissue along with a matched normal control sample. Patient 15 of the dataset. Sequenced in duplicate on HiSeq2000. |

#### Statistics and other methods

DeepTools2.0 [5] was used to assess genome-wide similarity of the sequencing replicates. All statistical tests were performed in R 3.6.1 (https://cran.r-project.org/mirrors.html). SPLICE-q's workflow figures were generated with Drawio (https://github.com/jgraph/drawio).

3. Dobin A, Davis CA, Schlesinger F, Drenkow J, Zaleski C, Jha S, et al. STAR: ultrafast

universal RNA-seq aligner. Bioinformatics. 2013;29:15–21. doi:10.1093/bioinformatics/bts635.
